# Neurophysiological alterations in the nucleus reuniens of a mouse model of Alzheimer’s disease

**DOI:** 10.1101/643023

**Authors:** D.A. Walsh, J.T. Brown, A.D. Randall

## Abstract

Transgenic mice that overproduce beta-amyloid (Aβ) peptides exhibit neurophysiological alterations at the cellular, synaptic and network levels. Recently, increased neuronal activity in nucleus reuniens (Re), has been linked to hyperexcitability within hippocampal-thalamo-cortical networks in the J20 mouse model of amyloidopathy. Here *in vitro* whole-cell patch clamp recordings were used to compare old pathology-bearing J20 mice and wild-type controls to examine whether alterations to the intrinsic electrophysiological properties of Re neurons could contribute to the amyloidopathy-associated Re hyperactivity. A greater proportion of Re neurons displayed a hyperpolarised membrane potential in J20 mice without changes to the incidence or frequency of spontaneous action potential (AP) generation. Passive membrane properties were independent of transgene expression. Re neurons recorded from J20 mice did not exhibit increased AP generation in response to depolarising current stimuli but did exhibit an increased propensity to rebound burst following hyperpolarising current stimuli. This increase in rebound firing does not appear to result from alterations to T-type calcium channels. Finally, in J20 mice there was an ∼8% reduction in spike width, similar to what we and others have reported in CA1 pyramidal neurons from multiple amyloidopathy mice. We conclude that alterations to the intrinsic properties of Re neurons may contribute to the hyperexcitability observed in hippocampal-thalmo-cortical circuits under pathological Aβ load.

**Key Points:**

- Alterations in the neurophysiology of hippocampal and cortical neurons has been linked to network hyperexcitability in mouse models of amyloidopathy.
- The nucleus reuniens (Re) is part of a cognitive network involving the hippocampal formation and prefrontal cortex. Increased cellular activity in Re has been linked to the generation of hippocampal-thalamo-cortical seizure activity in J20 mice.
- Re neurons display hyperpolarised resting membrane potentials in J20 mice. Passive membrane properties are unaffected by transgene expression. Re neurons recorded from J20 mice did not exhibit increased excitability in response to depolarising current stimuli but did exhibit an increased propensity to rebound burst following hyperpolarising current stimuli. This increased rebound firing was not a result of changes in T-type Ca^2+^ conductances. Finally we observed a decrease in AP width.
- These results help us understand how altered Re cellular neurophysiology may contribute to hippocampal-thalamo-cortical hyperexcitability in J20 mice.

## Introduction

The excess generation and/or reduced clearance of beta-amyloid (Aβ) peptides in the central nervous system (CNS) of patients with Alzheimer’s disease (AD) leads of the accumulation of amyloid plaques, a core pathological feature of the disease. The amyloid hypothesis of AD states that this imbalance between production and removal of Aβ within the CNS is a key determinant of the functional deficits, and accompanying neuronal loss, associated with AD (Hardy & Higgins, 1992; Selkoe & Hardy, 2016). The advent of mouse models of amyloidopathy has provided researchers with an incredibly powerful tool for studying the functional and behavioural abnormalities associated with an increased Aβ load in the CNS (McGowan *et al.*, 2006; Randall *et al.*, 2010). These transgenic models typically express causal genetic mutations associated with familial AD, resulting in overproduction of pathological forms of Aβ. One such model which is widely used to model amyloid pathology is J20 (PDGF-APPSw, Ind) mice (Mucke *et al.*, 2000). This transgenic line overexpresses human amyloid precursor protein (APP) with two mutations (Swedish and Indiana) under the platelet-derived growth factor (PDGF) promotor, resulting in increased Aβ production throughout the CNS. J20 mice recapitulate many of the hallmarks of AD pathology including amyloid plaque deposition (Wright *et al.*, 2013), synaptic loss (Hong *et al.*, 2016), cognitive impairment (Cheng *et al.*, 2007; Wright *et al.*, 2013), and network hyper-excitability (Palop & Mucke, 2009; Verret *et al.*, 2012; Hazra *et al.*, 2016).

The majority of studies investigating neurophysiological deficits in models of amyloidopathy have focused on altered synaptic function (Jacobsen *et al.*, 2006; Saganich *et al.*, 2006; Witton *et al.*, 2010) and associated functional abnormalities within defined neural networks, most commonly in the hippocampus and cortex. More recently, evidence has accumulated that alterations in the intrinsic properties of discrete neural populations appear to be a common phenotype arising from overexpression of Aβ, with changes consistently reported in intrinsic excitability profile and AP waveform (Brown *et al.*, 2011; Wykes *et al.*, 2012; Kerrigan *et al.*, 2014; Tamagnini *et al.*, 2015). With the assumption that neurophysiological changes at the level of single neurons will manifest themselves at the level of complex network activity underlying cognitive processing, these reports have provided important insight into the emergence of functional deficits characteristic of AD. However, as with work on synaptic function, such studies have focused almost exclusively on alterations to the intrinsic properties of neurons in the hippocampus or cerebral cortex.

In the last 25 years there has been a growing appreciation that specific thalamic nuclei, making up a portion of the limbic thalamus, are important at various levels of cognitive function (Aggleton & Brown, 1999; Aggleton, 2014). One of these the nucleus reuniens (Re), a midline thalamic nucleus, forms strong reciprocal connections with both the hippocampus and medial prefrontal cortex (Vertes *et al.*, 2015). Interest in Re has grown dramatically in recent years (Cassel *et al.*, 2013), with considerable evidence indicating a role for the Re in various forms of memory (Hembrook & Mair, 2011; Cholvin *et al.*, 2013; Hallock *et al.*, 2013; Pereira de Vasconcelos & Cassel, 2015) and the emergence of oscillatory synchrony between the HPC and mPFC during cognitive tasks (Hallock *et al.*, 2016; Roy *et al.*, 2017; Kafetzopoulos *et al.*, 2018). Spatial and goal-oriented neurons have been described in Re (Jankowski *et al.*, 2014, 2015; Ito *et al.*, 2015) and experimental inactivation of this area results in spatial deficits, reflecting those commonly observed in both AD patients and mouse models of AD. Recently a study, has linked increased c-fos immunoreactivity (>300%) in Re to cortical epileptiform activity frequently described in J20 mice (Hazra *et al.*, 2016).

In this initial observational study, we present data on the intrinsic membrane properties of Re neurons recorded from 12-14 month old J20 neurons and age-matched wild-type controls. To our knowledge, this is the first study which describes alteration in the intrinsic properties of Re neurons in a rodent model of dementia. The implications of these alterations in the context of aberrant network activity will be discussed.

## Methods

### Ethical approval

All work in this study was approved by the University of Exeter Animal Welfare Ethical Review Board. Animals were sacrificed in accordance with schedule 1 of the UK Animals (Scientific Procedures) Act 1986 and the subsequent amendments to the regulations of 2012, as implemented in response to directive 2010/63/EU of the European Parliament and of the Council on the protection of animals used for scientific purposes.

### Animals and tissue preparation

Male transgenic (TG) J20 mice (background strain: C57-Bl/6J) and wild-type (WT) littermate controls were bred in house at the University of Exeter. They were subsequently housed on a 12:12 light/dark cycle and granted *ab libitum* access to food and water. This study used mice of approximately 13 months of age (WT, mean 13.1 months, range 12.1-14.2 months, n = 12; TG, mean 13 months, range 12.4-14.3 months, n = 13). Following cervical dislocation, the brain was rapidly resected and placed within an ice-cold sucrose-based slicing medium consisting of (in mM): 189 Sucrose, 10 D-Glucose, 26 NaHCO_3_, 3 KCl, 5 Mg_2_SO_4_, 0.1 CaCl_2_, 1.25 NaH_2_PO_4_. Coronal sections of 300 μm thickness were sliced using a Leica VT1200 vibratome, and allowed to recover at room temperature for at least one hour prior to recording. Slice recovery was in our standard recording aCSF, composed of (in mM): 124 NaCl, 3 KCl, 24 NaHCO_3_, 1.25 NaH_2_PO_4_, 2 CaCl_2_, 1 MgSO_4_, 10 D-Glucose, gassed with carbogen (i.e. 95% O_2_ /5% CO_2_). A single coronal section containing the rostral Re was identified as described previously (Walsh *et al.*, 2017).

### Electrophysiological recordings

For recordings the slice containing rostral Re was transferred to a commercial submerged recording chamber (Warner Instruments), which was mounted on the stage of an upright microscope (Olympus BX51). The chamber was perfused with a continuous flow of temperature-controlled (32-33°C), carbogen bubbled, aCSF. Re neurons were visualised using infrared differential interference contrast optics and a CMOS USB 2.0 camera (ThorLabs). All recordings were made using the patch clamp technique. Pipettes (3-5 MΩ) were fabricated from borosilicate glass capillaries using a P-97 Flaming Brown micropipette puller. Pipettes were filled with either a potassium-gluconate based internal solution (for current-clamp recording) composed of (in mM): 140 K-gluconate, 10 NaCl, 10 HEPES free acid, 0.2 EGTA, 0.3 Na-GTP, 4 Mg-ATP, pH adjusted to 7.3 with KOH, or a cesium methanesulfonate based internal solution (voltage-clamp recording) composed of (in mM): 130 CsMeSO_4_, 20 NaCl, 10 HEPES free acid, 0.2 EGTA, 0.3 Na-GTP, 4 Mg-ATP, pH adjusted to 7.3 with CsOH. All recordings were made with a Multiclamp 700B amplifier (Molecular Devices) and digitised with a Digidata 1440A interface (Molecular Devices). All data were stored on a personal computer (Hewlett-Packard) using pClamp 10.4 software.

### Data analysis

Data were generally analysed using custom-written MATLAB scripts or, on occasion, using pClamp 10.4 software. A junction potential error of either −15 mV (K-Gluconate internal) or −9 mV (CsMeSO_4_ based internal) was corrected for arithmetically during data analysis. Statistical significance between genotypes was assessed using unpaired two-tailed Student’s t-tests, Mann-Whitney U tests, Kolmogorov–Smirnov tests or repeated-measure two-way analysis of variance (ANOVA) as appropriate, within the SPSS Statistics 22 software platform (IBM). Figures were prepared using Origin 2015.

## Results

Previous recordings made within our lab from young adult (16-18 weeks) male mice indicate that Re neurons typically display a relatively depolarised resting membrane potential (V_m_) and an associated propensity to fire spontaneous action potentials (APs) in the absence of any external depolarising input (Walsh *et al.*, 2017). To assess whether the spontaneous firing behaviours of Re neurons differed between WT and TG mice, V_m_ was recorded, in the absence of exogenous stimuli, for 60 s following breaking in to the whole cell mode. Commensurate with previous findings, the majority of Re neurons recorded from both WT and TG mice exhibit a depolarised (>-70 mV) membrane potential (WT, median −63.5 mV, TG, median −66.9 mV). However the proportion of rostral Re neurons exhibiting a relatively hyperpolarised V_m_ was increased in J20 mice when compared to WT controls (Fig 1A, WT, n = 43; TG, n = 49; p = 0.05, Kolmogorov–Smirnov test). As illustrated in figure 1A, approximately 40% of Re neurons recorded from TG mice displayed a V_m_ of ≤ −75 mV, compared to ∼10% of WT controls. Interestingly, this did not correspond to a significant decrease in the number of neurons exhibiting spontaneous AP generation in the absence of applied stimuli (Fig 1B, p = 0.25, Chi-squared test), with the majority of neurons of both genotypes firing at least 1 AP during the 60 s recording period.

**Figure 1.**
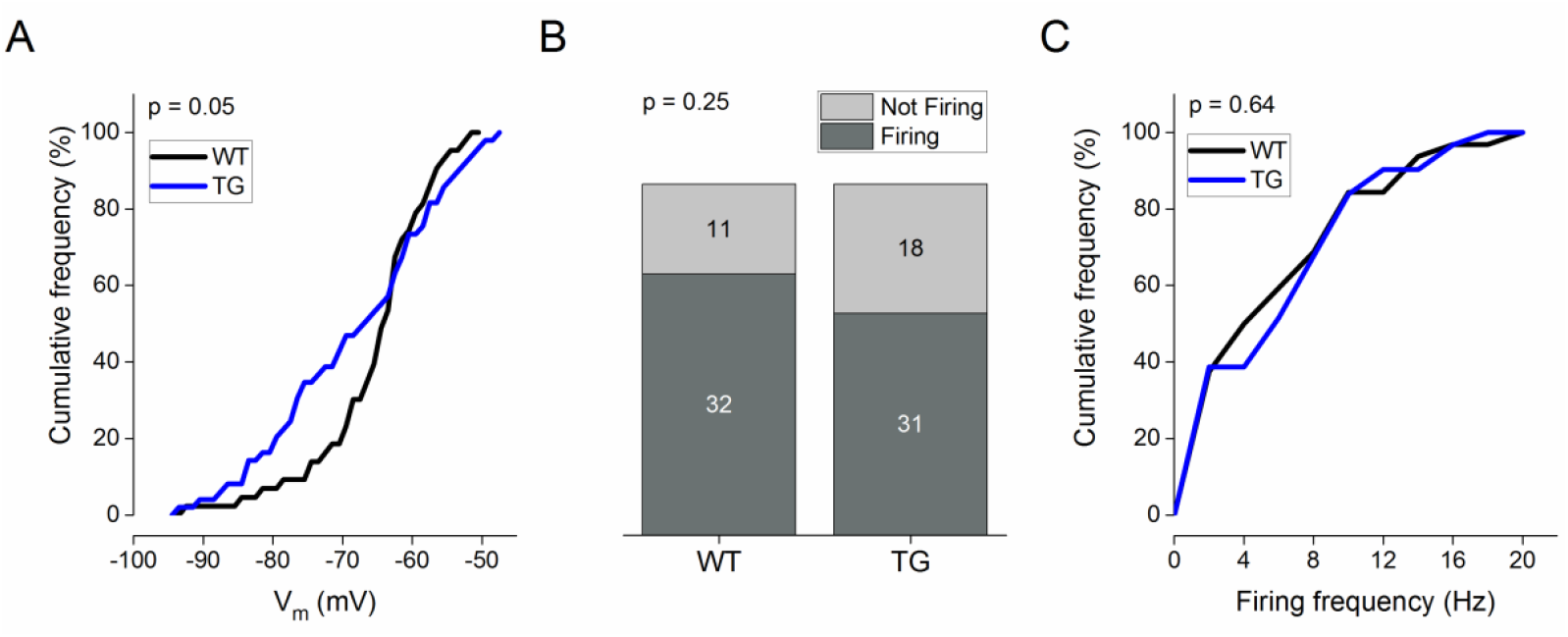
(A) A cumulative frequency plot displaying the distribution of V_m_ observed across groups. Black line represents WT neurons; blue line represents TG neurons. p value was calculated using a Kolmogorov–Smirnov test. (B) A cumulative column representation of the number of silent and spontaneously firing neurons across genotype. p value was calculated using a Chi-squared test. (C) Plot showing, for firing cells, the mean firing frequency for each genotype.

We have previously described 4 different populations of Re neurons based on spontaneous firing behaviours (Walsh *et al.*, 2017) and found, predictably, that V_m_ varies between these populations. As such, the increased proportion of hyperpolarised neurons in J20 mice could be a manifestation of an increase in the relative proportion of active neurons displaying low firing frequencies associated with hyperpolarised V_m_. To test this we examined the population distributions of spontaneous neuronal firing frequencies. Firing frequency was calculated by dividing the number of APs observed within the 60 s recording by 60. No difference was observed between WT and TG mice (Fig 1C, WT, n = 32; TG, n = 31, p = 0.64, Kolmogorov–Smirnov test). Another possible explanation for the change in V_m_ without a parallel change in either the proportion of cells exhibiting APs or the frequency of spontaneous APs is a change in the V_m_ of neurons which generate no spontaneous APs. However, the V_m_ of quiescent neurons was not significantly different between WT and TG neurons (WT, mean −74.4 ± 2.8 mV, n = 11; TG, mean −76.3 ± 2.2 mV, n = 18; p = 0.59, unpaired, two tailed student’s t-test, data not shown).

In order to compare passive membrane properties between Re neurons recorded from WT and TG mice, a 500 ms, 30 pA hyperpolarising current pulse was passed across the neuronal membrane. For such assessments recordings were made from a set pre-stimulus membrane potential of −80 mV, achieved using a steady state bias current injection. This was to prevent the cell to cell variability in resting membrane potential from introducing variability in intrinsic properties which, as in all neurons, express some degree of voltage-dependence. There were no statistically significant differences observed when comparing the passive properties between WT and TG Re neurons. These cells exhibit no Ih mediated sag (Walsh et al 2017) hence the input resistance (R_i_) was calculated from Ohm’s law by dividing the mean voltage deflection calculated during the last 50 ms of the −30 pA hyperpolarising sweep by the amplitude of the negative current injection. Median R_i_ for WT neurons was 426 MΩ compared to 431 MΩ for TG neurons (Fig 2A, p = 0.88, Mann-Whitney U test). The membrane time constant (τ) was calculated by fitting a single exponential curve to the charging trajectory of the membrane potential between 20-80% of the peak amplitude. The median τ was 29.6 ms for WT neurons as compared to 29.1 ms for TG neurons (Fig 5.4B, p = 0.35, Mann-Whitney U test). Membrane capacitance was approximated by calculating the ratio of τ/R_i_. The median capacitance for WT neurons was 64.0 pF as compared to 54.8 pF for TG neurons (Fig 5.4C, p = 0.54, Mann-Whitney U test).

**Figure 2.**
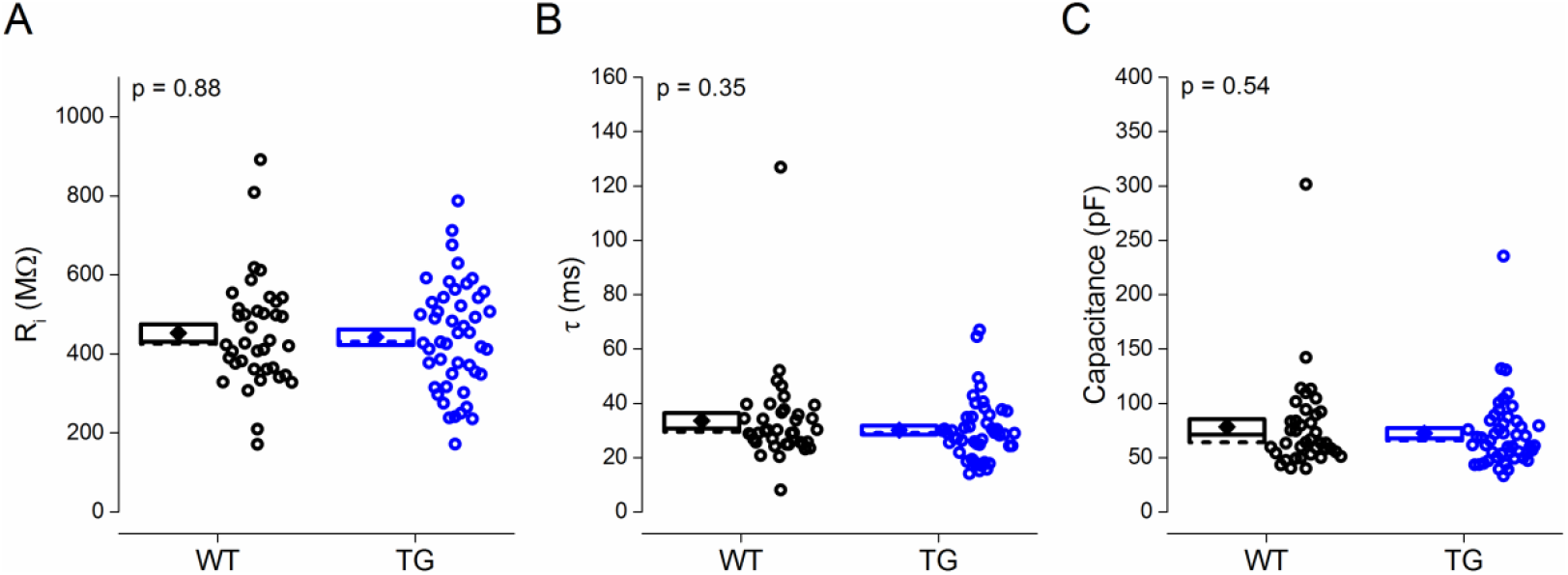
Passive membrane properties were independent of transgene expression. (A-B) Plots of (A) R_i_ and (B) τ, calculated from a 500 ms, 30 pA hyperpolarising current injection. (C) Plot showing capacitance calculated as τ/R_i._ Diamond represents mean, dashed line represents median, and box represents SEM. All p values were calculated using a Mann-Whitney U test

**Figure 3.**
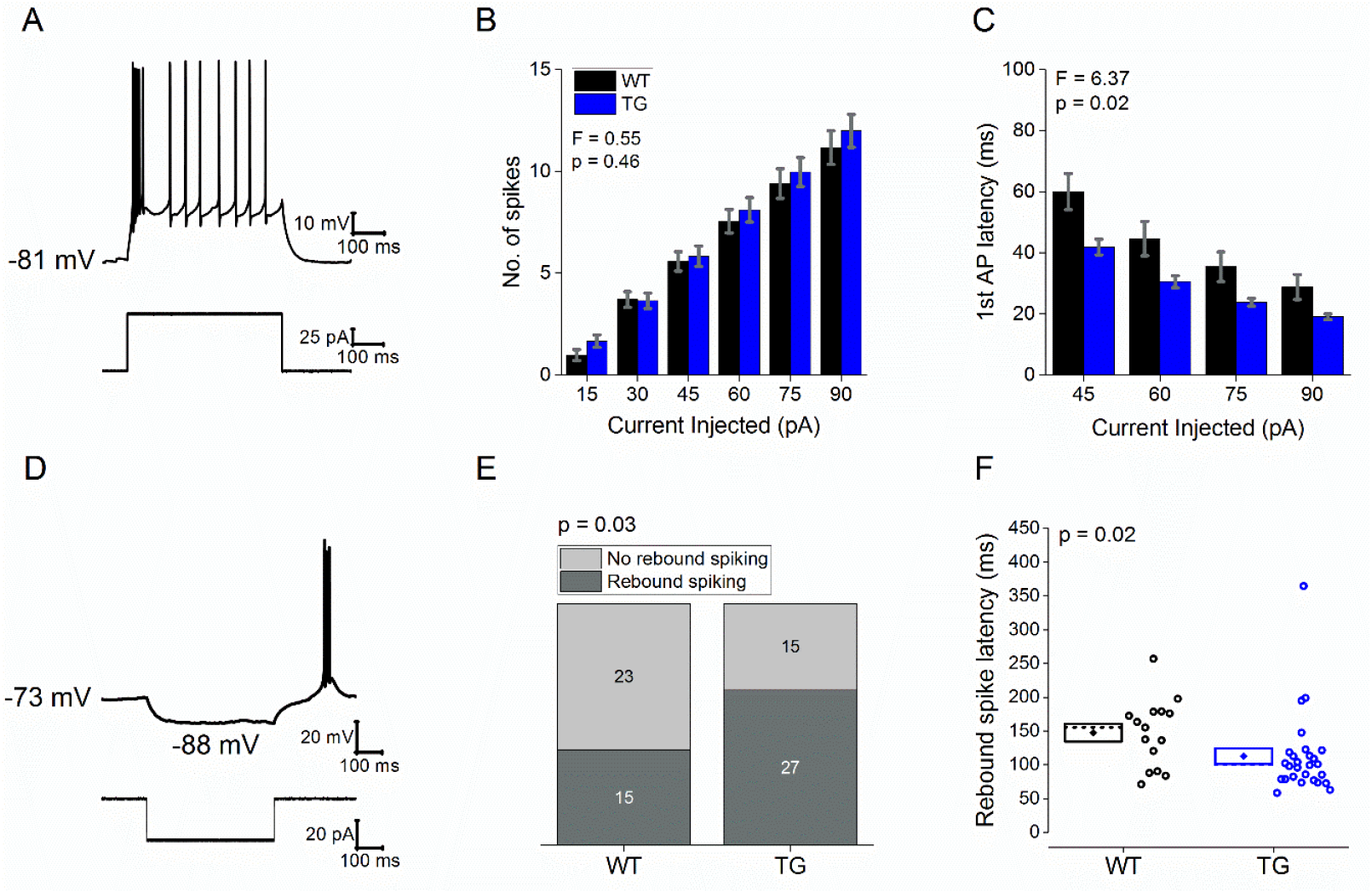
(A) Sample voltage response (top) and current trace (bottom) of a 90 pA, 500 ms depolarising current injection from a prestimulus potential of −80 mV. Plots showing (B) the mean number of APs and (C) mean latency to the first AP, produced in response to a series of 500 ms depolarising current injections from a prestimulus potential of −80 mV. F and p values calculated using a 2 way RM-ANOVA are shown. (D) Sample voltage (top) and current (bottom) trace of a −30 pA, 500 ms hyperpolarising current step from a set membrane potential of −73 mV. (E) A cumulative column representation of the number of neurons which did not exhibit rebound firing and the number of neurons which exhibited rebound firing from a holding potential of −73 mV. p values were calculated using a Chi-squared test. (F) Plot showing the latency to the peak of the 1^st^ rebound spiken. Diamond represents mean, dashed line represents median, and box represents SEM. p value was calculated using a Mann-Whitney U test.

**Figure 4.**
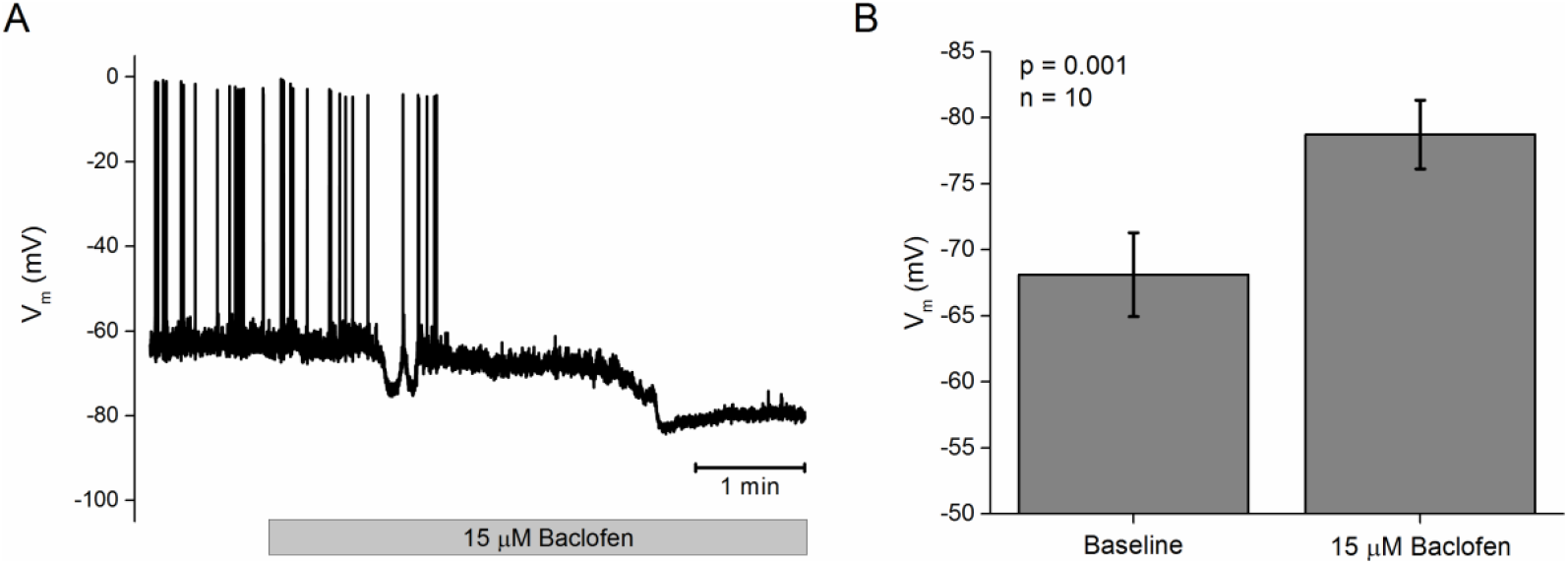
(A) Sample Voltage trace showing the effect of 15 μM Baclofen on Vm. (B) Bar graph showing the mean effect of Baclofen on Vm. Error bars represent sem. p value was calculated using a paired two tailed Student’s t-test.

**Figure 5.**
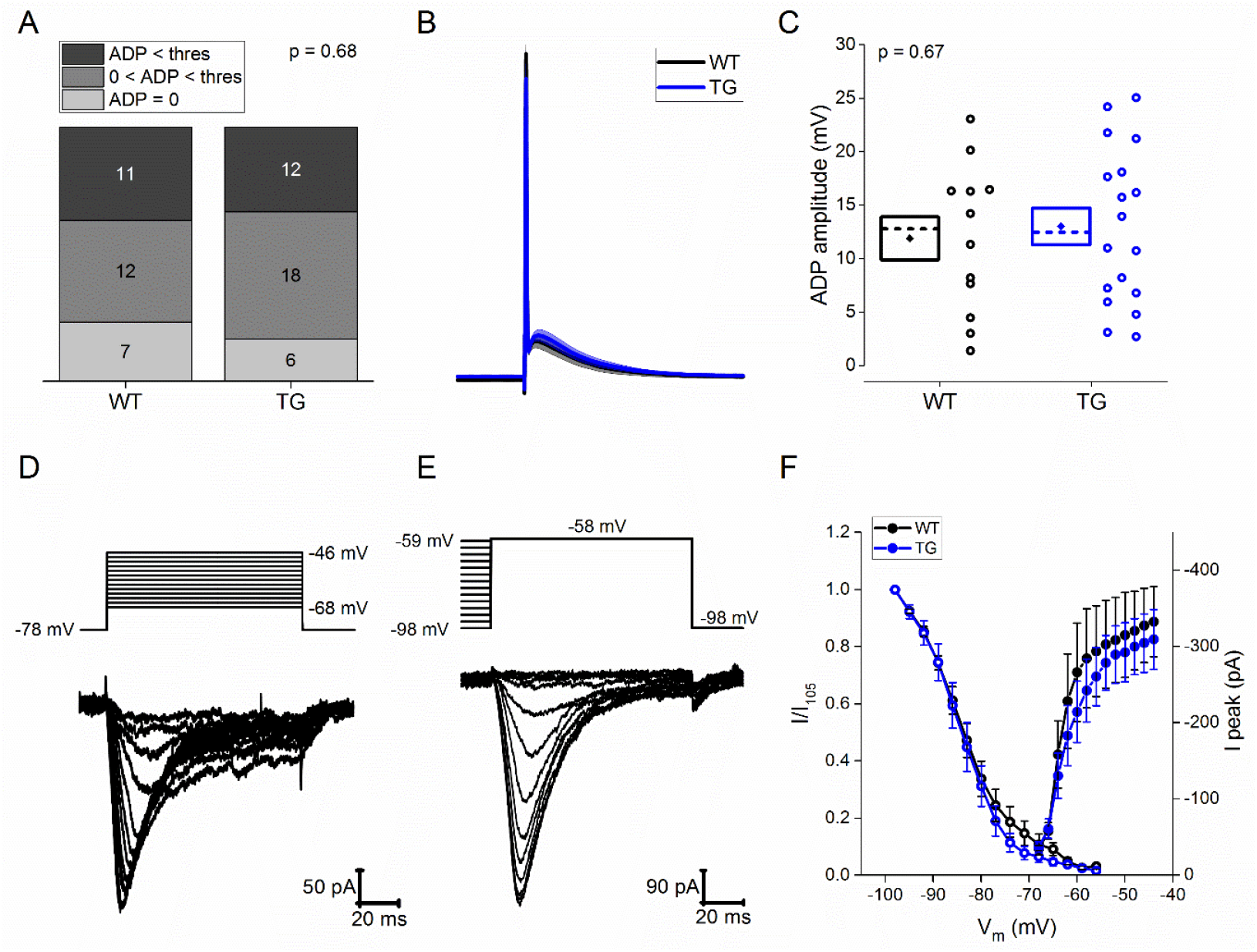
(A) Cumulative column representation of the number of neurons which do not exhibit an ADP, the number of neurons exhibiting a subthreshold ADP and the number of neurons which exhibit a suprathreshold ADP across genotype. This was measured from a holding potential of −80 mV with a 1.25 ms 2 nA current stimulus. p value was calculated using a Chi-squared test. (B) Average voltage trace in response to a short (1.25 ms), large (2 nA) current injection in cells exhibiting a subthreshold ADP. Line represents mean and shaded area represents SEM. (C) Plot showing the ADP amplitude in neurons exhibiting a subthreshold ADP. Diamond represents mean, dashed line represents median, and box represents SEM. p value was calculated using an unpaired, two tailed student’s t-test. (D) Sample voltage trace and the subsequent current response to a series of 2 mV incremental depolarising voltage steps (from −75 – −47 mV) from a prestimulus potential of −85 mV using a CsMeSO_4_ based internal solution. (E) Sample voltage trace and the subsequent current responses to a series of depolarising voltage steps to −65 mV from an incrementally depolarised prestimulus potential (−105 mV – −66 mV) were used to calculate the steady state inactivation curve of T-type Ca^2+^ channels. (F) Average I-V plot (closed circles) of the peak current observed in response to a series of 2 mV voltage steps. Inactivation curve (open circles) showing the average voltage at which T-type Ca^2+^ channels inactivate.

To compare the excitability of Re neurons between WT and TG mice, a series of incremental depolarising current injections were made ranging in amplitude from 15 to 90 pA. These were also applied at a fixed pre-stimulus membrane potential of −80 mV. A sample trace showing the response to a 90 pA current injection is displayed in Figure 3A. The mean number of APs and the latency to the first AP produced in response to the series of current injections were used to quantify excitability. The mean number of APs elicited, calculated for each population by dividing the sum of APs (including zero spike sweeps) produced by population cell count, was not affected by transgene expression (Fig 3B, WT, n = 38; TG, n = 47; F = 0.55, p = 0.46, 2 way RM-ANOVA). Latency to the first AP was measured in response to the four largest current injections (45 – 90 pA), as each resulted in at least one AP in the vast majority of Re neurons in both WT (97%) and TG (95%) mice. Latency was calculated as time taken from the initiation of the depolarising current stimulus to the peak of the first AP. The latency to the first AP was ∼32% shorter in Re neurons recorded from J20 mice (Fig 3C, WT n = 37; TG, n = 45; F = 6.37, p = 0.02, 2 way RM-ANOVA). From more depolarised membrane potentials, a sizeable proportion of Re neurons display rebound firing following a 500 ms hyperpolarising current injection. This appears to result largely from the de-inactivation of T-type Ca^2+^ channels afforded by hyperpolarization (Walsh *et al.*, 2017). To assess whether the propensity for rebound firing was genotype-associated expression, a 500 ms, 30 pA hyperpolarising current injection was injected from a set pre-stimulus potential of – 72 mV (Fig 3D), which sits between the overall average resting membrane potential and the mean resting potential of the subgroup of cells that did not exhibit spontaneous firing. TG neurons had a higher propensity to rebound fire than WT neurons (Fig 3E, WT, 39%; TG, 64%; p = 0.03, Chi-squared test). For the cells which exhibited such hyperpolarization induced firing, rebound latency, defined as the time taken from the cessation of the current stimulus to the peak of the first rebound spike, was also calculated. This was significantly shorter in TG (median 112.5 ms) as compared to WT (median 147.4 ms) neurons (Fig 5.3D, WT, n = 15; TG, n = 27; p = 0.02, Mann-Whitney U test).

GABA_B_ receptors are widely expressed throughout the mouse thalamus. When activated at synapses the resultant post-synaptic hyperpolarization can reliably generate rebound burst firing in thalamic relay neurons (Ulrich *et al.*, 2018). A radioligand binding analysis reported many baclofen sensitive GABA binding sites in Re (Hosford *et al.*, 1995), and the same study provided evidence of directionally opposed outcomes of GABAB receptor agonism and antagonism on spike and wave discharges in the lethargic mouse model of absence seizures. Commensurate with this, the Allen Brain Atlas (http://mouse.brain-map.org) indicates that both the subunits required to make the GABA_B_ receptor are robustly expressed in Re. However to date, a comprehensive characterisation of the synaptic receptors present on Re neurons has not been published. Hence, to confirm a likely neurophysiological role for GABA_B_ receptors in Re neurons, we recorded neurons from ∼8 month old WT mice and applied 15 μM of GABA_B_ receptor agonist R-Baclofen (the active enantiomer) via the aCSF perfusing the recording chamber. Following break in to the whole cell mode, Vm was recorded for at least one minute to establish a baseline. Baclofen was then washed on and the cellular response was followed for 5 minutes. Baclofen produced a hyperpolarisation in Re neurons, comparing baseline (mean −68 mV) and the fifth minute of application (−78 mV) (Fig 4B, n = 10, p = 0.001, paired, two tailed students t-test), suggesting that Re neurons have the potential to display rebound firing in response to GABAB-receptor mediated inhibitory synaptic input. Exogenous baclofen also reduced the variance in the membrane potential likely as a result of reduced presynaptic release of glutamate and/or GABA at synapses made on to Re cells.

As previously mentioned, T-type calcium currents are necessary for rebound firing in Re neurons. We wished to assess whether the increased propensity to rebound fire in TG mice may result from of alterations to the amplitude or kinetics of T-type calcium channels. One would expect that such a change would manifest as an alteration in the number of neurons exhibiting sub-threshold and supra-threshold afterdeoplarisations (ADP), a feature underpinned in Re neurons by T-type calcium conductances (Walsh *et al.*, 2017). In order to quantify ADP amplitude a single spike was elicited with a large (2 nA), short (1.3 ms) depolarising current injection applied at a set prestimulus potential of – 80 mV. Neurons were divided into one of three groups qualitatively; in any neuron which lacked a clear depolarising phase following the peak of AP the ADP was interpreted to be zero. Any neuron where the amplitude of the ADP was not sufficiently large to elicit an AP was defined as exhibiting a sub-threshold ADP, while if the ADP was of a great enough amplitude to cross AP threshold it was defined as supra-threshold. The proportion of neurons exhibiting supra-threshold, sub-threshold, or no quantifiable ADP was similar between WT and TG Re neurons (Fig 5A, p = 0.68, Chi-squared test). Direct quantification of ADP amplitude was restricted to neurons displaying sub-threshold ADPs, as the presence of one or more ADP-driven APs obstructs faithful measurement in the suprathreshold population. An average trace of the V_m_ in response to the depolarising current injection in neurons displaying a sub-threshold ADP is displayed in Figure 5B. ADP amplitude did not significantly differ between WT (mean 11.9 ± 2.0 mV) and TG (mean 13.0 ± 1.7 mV) mice (Fig 5C, WT, n = 12; TG, n = 18; p = 0.67, unpaired, two tailed students t-test).

In order to confirm that the activity of T-type calcium channels was unaltered in J20 mice, we carried out a series of voltage-clamp recordings using a CsMeSO_4_ based internal solution to directly measure the amplitude and voltage-gated kinetics of the low threshold Ca^2+^ current directly. A series of voltage steps was applied, ranging from −68 to −40 mV, from a holding potential of −78 mV (see Figure 5D). In response to sufficiently large voltage steps an additional inward current was observed which was evidently larger and displayed longer activation and inactivation kinetics than the inactivating T-type channel currents (Walsh *et al.*, 2017). These likely arise from the initial activation of a large HVA Ca^2+^ channel current component. In an attempt to minimise the confounds of measuring the amplitude of T-type Ca^2+^ current, analysis was restricted to those voltage steps ranging from −75 mV to −46 mV, where HVA Ca^2+^ current activation was minimal.

The steady state inactivation profile of the channel was studied using a depolarising voltage step to – 58 mV from a pre-step potential incrementally increasing from −98 mV to −59 mV. A representative recording of the evoked currents is displayed in Figure 5E. Average I/V plots of maximal inward current and inactivation curves are displayed in Figure 5F. Visual examination of the curves indicated that there is no clear hyperpolarising or depolarising shift in either plot. In light of the data presented in Figures 4, one can say with a degree of confidence that alterations the amplitude or voltage-gated kinetics of T-type Ca^2+^ channels do not play a causal role in the increased propensity of Re neurons to rebound fire.

Studies from our lab (and others), have reliably shown that the AP width of CA1 pyramidal neurons is approximately 10-15% narrower in mouse models of amyloidopathy (Brown *et al.*, 2011; Wykes *et al.*, 2012; Kerrigan *et al.*, 2014; Tamagnini *et al.*, 2015). This is something we have also observed in CA1 cells of J20 mice, CA1 cells in TAS-TPM mice and CA1 pyramids of rTg4510 tauopathy mice (Francesco Tamagnini, unpublished observations). To assess if a similar alteration is present in Re neurons we compared the AP waveform properties of WT and TG mice. AP waveform properties were measured from the first spike generated in response to the 90 pA, 500 ms current injection from a pre-stimulus potential of −80 mV (see Fig 3A). The average peak aligned AP waveform is for WT and TG neurons is displayed in Figure 6A, with corresponding phase plot shown in Figure 6D. AP peak was measured as the absolute zenith of the action potential waveform. There was no difference in the AP peak between WT (mean 18.4 ± 1.3 mV) and TG (mean 17.2 ± 1.1 mV) mice (Fig 6B, WT, n = 38; TG, n = 47; p = 0.49, unpaired, two tailed Student’s t-test). AP half-width was measured as AP width at half height of the AP, where height was defined as the voltage difference between AP peak and AP threshold. AP width was approximately 8% shorter in Re neurons recorded from TG mice (Fig 6C, WT, mean 0.68 ± 0.02 ms, n = 38; TG, mean 0.63 ± 0.01 ms, n = 47; p < 0.01, unpaired, two tailed student’s t-test). AP threshold was defined as the voltage at which the first derivative of the membrane voltage (dV/dt) during the AP waveform exceeded 15 mV/ms. AP threshold did not significantly differ between WT (median −52.1 mV) and TG (median −53.1 mV) mice (Fig 6E, WT, n = 38; TG, n = 47; p = 0.24, Mann-Whitney U test). Maximal rate of rise (max dV/dt) was defined as the peak value of the first derivative of the AP waveform. Max dV/dt did not significantly differ between WT (median 296 mV/ms) and TG (median 317 mV/ms) mice (Fig 6F, WT, n = 38; TG, n = 47; p = 0.16, Mann-Whitney U test).

**Figure 6.**
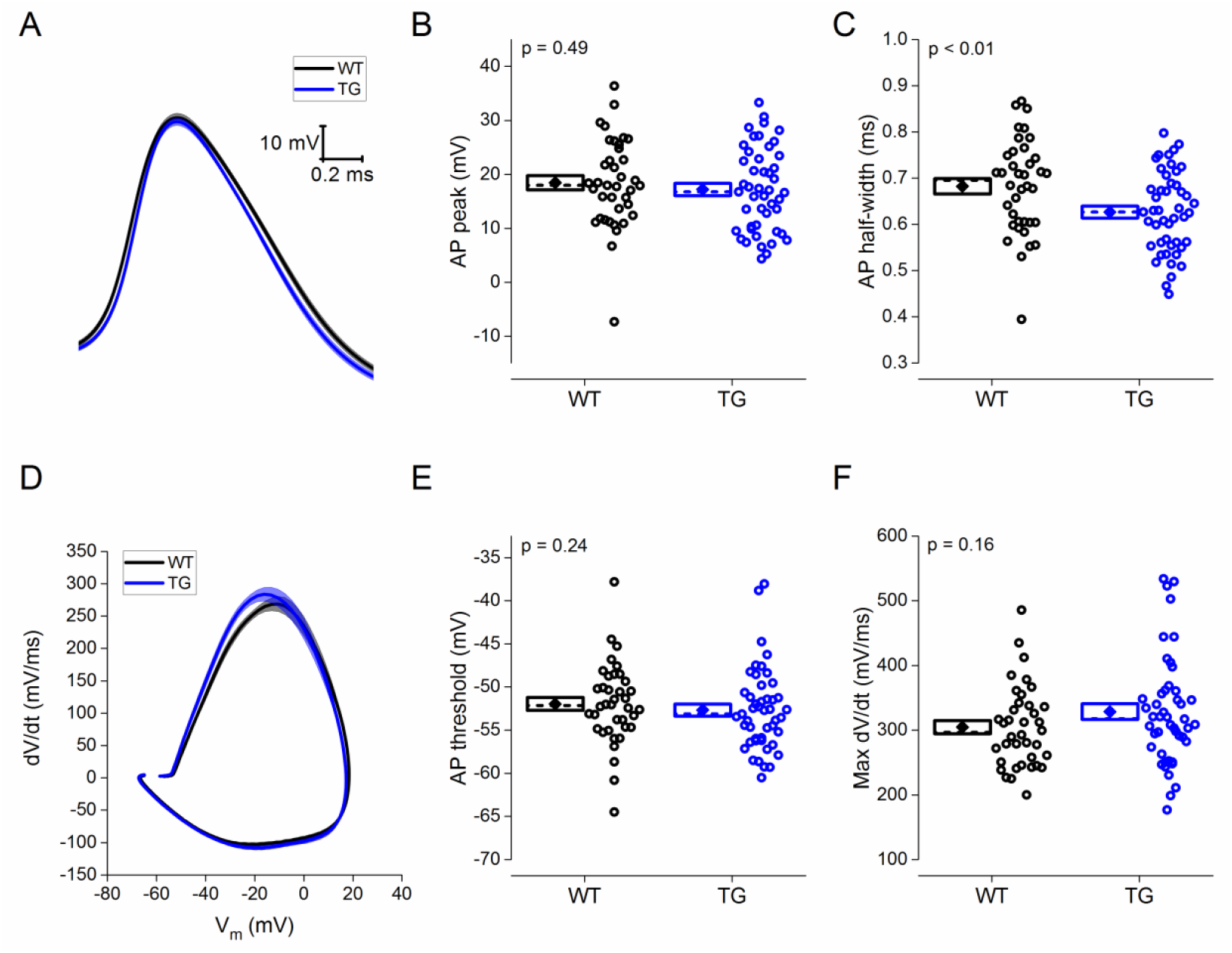
(A) Average waveform of the first AP generated in response to 90 pA, 500 ms depolarising current injection. Plots showing (B) the absolute peak voltage value of the AP and (C) the AP width at half height. Diamonds represent means, dashed lines represent medians, and the box outline represents SEM. All p values were calculated using an unpaired, two tailed student’s t-test. (D) Average phase plot of first generated AP, plotting the first derivative of the AP voltage against AP voltage. Plots showing (E) the AP threshold and (F) the maximal rate of AP rise. Diamonds represent means, dashed lines represent medians, and the box outline represents SEM. All p values were calculated using a Mann Whitney U test.

## Discussion

Re is integral to normal cognitive processing. This is proposed to be largely underpinned by its influential role within a cognitive network containing the hippocampus, the mPFC and possibly also the subiculum. This study in the widely used J20 mouse model of amyloidopathy provides the first report of alteration to Re electrophysiology in a mouse model of dementia. Alterations to the intrinsic membrane properties of Re neurons are likely have important consequences for learning and memory. This study indicates that the likelihood that Re neurons will produce burst firing following hyperpolarization (Figure 3), and the AP waveform (Figure 6) are altered in face of an amyloid pathology. A shortening in the latency to fire for a given depolarizing stimulus was also noted in the disease model, which is a likely indication of somewhat heightened excitability (Figure 3) and could also change spike timings within reciprocally connected networks.

These novel findings indicate that, in addition to synaptic and network dysfunction previously identified in brain regions more typically associated with memory (e.g. HPC), alterations to the intrinsic cellular electrophysiological properties of Re neurons may contribute to the well-characterised cognitive deficits exhibited by J20 mice (Cheng *et al.*, 2007; Verret *et al.*, 2012), and by extension potentially those experienced by sufferers of AD. Alterations to intrinsic properties of a neuronal population by definition arise downstream from changes to the expression/activity of the myriad voltage-gated ion channels present on neuronal membranes. Our own work (Walsh *et al.*, 2017) indicates that Re neurons appear to differ in significant ways from the much more widely studied relay neurons within sensory aspects of the thalamus. For example, they have a higher input resistance and seem to entirely lack HCN channels and a resulting “sag” potential they generate. Although a start has been made, a comprehensive understanding of the ionic conductances that shape various aspects of the neurophysiology of Re neurons is lacking. Analysis of the channelome in a single cell-level RNA sequencing analysis would be instructive in this regard, and an additional comparative dataset from age-matched J20 mice may help shed light on the neurophysiological outcomes outlined here. However, in the absence of these molecular and associated electrophysiological data, this study confines itself to providing a first descriptive account of intrinsic alterations to Re neurons in J20 mice.

### Increased propensity of Re neurons to burst fire

Potentially the most impactful finding of this study is the finding that Re neurons in J20 mice may display enhanced burst firing *in vivo* in response to both depolarising and hyperpolarising stimuli. As previously described (Walsh *et al.*, 2017), most Re neurons in adult brain slices typically display a relatively depolarised V_m_ and consequent propensity to tonically fire spontaneous APS in the theta frequency range. Similarly to typical thalamic relay nuclei, the membrane potential of Re neurons dictates whether they display tonic or burst firing when depolarised. At more hyperpolarised resting potentials, T-type calcium channels are available to activate in response to sufficient depolarisation resulting in a high-frequency burst of action potentials, which can be as fast as 300 Hz. Conversely, at more depolarised resting potentials, T-type calcium channels reside in inactivated states resulting in relatively low frequency tonic firing in response to depolarising stimuli. In J20 mice, a greater proportion of Re neurons exhibited a hyperpolarised membrane potential than WT controls and consequently, the proportion of Re neurons predisposed to high frequency burst firing (=< −80 mV) is greater in the disease model.

Similarly, Re neurons (along with typical thalmic relay neurons) can readily display bursting behaviour following a prolonged (500 ms – 1 s) hyperpolarising current stimulus. This results from the recovery from inactivation (i.e. deinactivation) of T-type calcium channels during the hyperpolarising step. The increased propensity of Re neurons to display rebound spiking in response to hyperpolarising current injections indicates Re neurons are also highly likely to display increased rates of burst firing in response to hyperpolarising stimuli *in vivo*. But what hyperpolarizing influence might deinactivate Re neurons LVA Ca^2+^ channels in vivo and lead to such high frequency rebound burst firing? Re neurons unquestionably lack the long-lasting, post-burst, Ca^2+^-dependent, afterhypolarizations exhibited by some other CNS neurons; indeed they generally exhibit marked post burst ADPs instead. Consequently, we propose the most likely source of sufficiently prolonged and large hyperpolarisations of Re neurons is synaptic activation of postsynaptic GABA_B_ receptors. Re neurons certainly receive GABAergic inputs, for example, at room temperature, spontaneous miniature GABA_A_ receptor-mediated IPSCs have been recorded at circa 2 Hz under recording conditions (i.e. intracellular Cs^+^) where GABA_B_ receptors would be blocked (Xu & Südhof, 2013). In our recordings from cells at physiological temperature, using K^+^-containing pipette solutions, Re neurones exhibit spontaneous GABAergic iPSCs at approximately 10 Hz in the absence of TTX. Indeed, the impact of inhibitory synaptic drive is exemplified by significant relationship between chloride equilibrium potential and the resting potential of Re neurons (DW and AR, unpublished observations). Furthermore, a manipulation (local virally-mediated Neuroligin 2 knockdown) that reduces GABAergic drive to Re has significant behavioural outcomes (Xu and Sudhof, 2013). We are unaware of any direct neurophysiological demonstration of GABA_B_-mediated IPSPs in Re neurons, although our data (Figure 4) indicate the required coupling of GABA_B_ receptors to K^+^ channels appears to be present. It would be interesting to develop a method to evoke monosynaptic IPSPs in Re neurons in slices, perhaps using an optogenetic strategy.

A number of lines of evidence support the assertion that the output mode (tonic vs burst firing) of Re neurons can profoundly influence both hippocampal and cortical activity. Burst firing in thalamic relay nuclei in response to hyperpolarising current is associated with both thalama-cortical delta oscillations, commonly seen during slow wave sleep, and spike and wave discharges (SWDs) observed during absence seizures. Notably increasing the power of Re delta oscillations by infusion of NMDA antagonist ketamine can impose delta oscillations onto the hippocampus. Similarly direct optogenetic activation of Re at delta frequency can impose an increase in delta power upon the HPC, resulting in cognitive deficits (Duan *et al.*, 2015). Meanwhile in a mouse model of atypical absence epilepsy in which GABA_B_ receptors are overexpressed post-natally, the emergence of SWDs within a cortico-thalamo-hippocampal circuit and associated cognitive deficits are dependant of the activity of GABA_B_ receptors in the midline thalamus (including Re) (Hosford *et al.*, 1995; Wang *et al.*, 2009).

Recently, a study which focused on the contribution of the thalmo-cortical system to seizure activity in J20 mice identified a 3-4 fold increase in spontaneous Re activity. Since both cognitive deficits and non-convulsive seizures (with associated SWDs) are reliable phenotypes reported in J20 mice, this present study raises the intriguing possibility that it is an increase in the propensity of Re neurons to burst fire that plays a causal role in the memory impairments and seizure activity seen in J20 mice. It would be of significant interest to perform longitudinal in vivo recordings of single unit activity in Re of J20 mice. Deep brain stimulation (DBS) has previously been used to reduce regional rebound bursting to great therapeutic effect in Parkinson’s disease (Meijer *et al.*, 2011; Cury *et al.*, 2017). This raises the intriguing possibility that DBS of Re could be a viable therapeutic strategy in the treatment of AD. Support for this notion is provided by a study by Arrieta-Cruz *et al.* (2010) who showed that 25 Hz deep brain stimulation of the midline thalamus facilitated acquisition of object recognition memory in the TgCRND8 mouse model of amyloidopathy.

### Excitability

With Re neurons preset to −80 mV no significant changes to the passive membrane properties or number of spikes elicited in response to a series of depolarising current injections in J20 mice were observed. There was, however, a ∼30% decrease in the latency to the 1^st^ spike both following initiation of depolarising stimuli and cessation of a hyperpolarising stimulus. As the predominant firing frequency observed over the entire course of a 500 ms depolarising stimulus in Re neurons is approximately 20 Hz, the 10 – 20 ms decrease in first spike latency is unlikely to manifest as an appreciable increase in spiking rate in response to prolonged depolarisation. This raises the interesting question as to whether the decrease in spike latency reported is likely to alter, in some fundamental sense, the information Re neurons transmit within cognitive networks. Notably such a change in latency could be more impactful when the source of depolarization is in the form of the transient current flux underpinning the depolarizing envelope of an EPSP rather than a “square-wave” current stimuli most typically used in experimental work. Firstly, this could result in a greater likelihood of a spike occurring at all for near threshold synaptic events, whereas for more robust excitatory inputs spike timing could be altered. Certainly whether spike timing or gross spike rate represent the fundamental unit of neural computation in the CNS is still the subject of debate within the neuroscientific community (London *et al.*, 2010; Bruno, 2011; Brette, 2015).

The more immediate question to our eyes is whether alterations to the excitability profile of Re neurons contribute to the increased risk of seizure activity commonly observed following prolonged exposure to increased Aβ load in both mouse and human. A primary motivation in the study of neuronal excitability in mouse models of AD, certainly within our lab, has been to try and link alterations at a neuronal level to hyperexcitability observed in hippocampal and cortical networks. Re neurons appear hyperexcitable in J20 mice and studies show that excess excitation of Re neurons results in increased propensity of convulsive seizures (Hirayasu & Wada, 1992; Luna-Munguia *et al.*, 2017). However the evidence presented in this work does not support the view that Re hyper-excitability in response to depolarising stimuli contributes to the increased seizure rates in J20 mice.

### Alterations in AP waveform

This study also uncovers the novel finding that the AP waveform of Re neurons is significantly narrower in J20 mice. The approximate 8% reduction in Re spike width is reminiscent of the spike narrowing of hippocampal CA1 pyramidal neurons in other mouse models of amyloidopathy, including PSAPP, PDAPP, CRND8 (Brown *et al.*, 2011; Wykes *et al.*, 2012; Kerrigan *et al.*, 2014; Tamagnini *et al.*, 2015), and indeed J20 mice (F. Tamagninini and AR, unpublished data). This finding was unexpected, as thalamic neurons differ substantially from CA1 pyramidal neurons in their expression profile of voltage-gated ion channels. The concordance in these findings both across models of amyloidopathy within CA1, and now across CA1 and Re in J20 mice suggests that the narrowing of APs is a reliable and potentially widespread phenomenon across the CNS in response to excess Aβ levels. Indeed we have also seen a similar change in some hippocampal GABAergic interneurons (Francesco Tamagnini and AR, unpublished observations).

Following initiation of an AP, a plethora of voltage-gated Na^+^, K^+^ and Ca^2+^ are recruited. Given the kinetics of the voltage-gated Ca^2+^ channels activated, Ca^2+^ flux across the membrane increases in an exponential fashion over the duration of the waveform. Indeed mathematical modelling of experimental data has suggested that a similarly modest decrease in spike width (10-15%) in CA1 neurons can lead to up to a 40% reduction in Ca^2+^ entry during an AP (Kerrigan *et al.*, 2014). Ca^2+^ ions are vital for an array of normal cellular processes and a decrease in Ca^2+^ entry would undoubtedly have marked effects in normal cellular functioning. For example, given the assumption that APs recorded at the soma reflect the AP waveforms that subsequently arrive in presynaptic terminals, reduced Ca^2+^ flux resulting from a narrower AP would lead to a reduction in the probability of neurotransmitter release. This will likely reduce the ability of Re to influence the activity of its various downstream targets including hippocampal area CA1 and the mPFC. For example, although studies focusing on alterations in basal synaptic transmission to CA1 in J20 have focused exclusively on the Schaffer Collateral (CA3 → CA1) pathway, a reduction in Re neurotransmitter release would likely manifest itself as a reduction in the amplitude of a field EPSP recorded following stimulation of presumed temporoammonic (EC → CA1) fibres in acute hippocampal slices. Such a finding would could interpreted falsely as an alteration to entorhinal cortex input to CA1.

### Limitations of in vitro recordings

Recordings made in brain slices are both high-throughput and convenient when compared to comparable in vivo alternatives. As such, we feel that this preparation is perfectly suited to this initial characterisation of alterations to Re intrinsic neurophysiological properties in a mouse model of dementia. However caution should be taken before extrapolating results viewed in vitro, where many inter-region connections are severed, to expected activity within an intact functioning neuronal network. This is especially true in thalamic neurons whose V_m_ (and associated functional output) is highly malleable in the face of excitatory and inhibitory modulatory input (Sherman & Guillery, 2006). For example, recent evidence suggests activation of mPFC-Re connections modulates burst firing in Re (Zimmerman & Grace, 2018). An important next step will be identifying which of the changes presented in this work are also evident in vivo. Importantly, doing so would allow these alterations to be correlated directly with behavioural deficits. In vivo, comparable alterations would likely have functional consequences within cognitive networks as an ever building literature suggests that the activity of Re can profoundly influence both hippocampal and cortical activity (Hosford *et al.*, 1995; Drexel *et al.*, 2011; Zhang *et al.*, 2012; Duan *et al.*, 2015).

